# Highly reduced complementary genomes of dual bacterial symbionts in the mulberry psyllid *Anomoneura mori*

**DOI:** 10.1101/2024.05.10.593479

**Authors:** Yuka Yasuda, Hiromitsu Inoue, Yuu Hirose, Atsushi Nakabachi

## Abstract

The genomes of obligately host-restricted bacteria suffer from accumulating mildly deleterious mutations, resulting in a drastic size reduction. Psyllids (Hemiptera) are phloem sap-sucking insects with a specialized organ called the bacteriome, which typically harbors two vertically transmitted bacterial symbionts: the primary symbiont “*Candidatus* Carsonella ruddii” (*Gammaproteobacteria*) and a secondary symbiont phylogenetically diverse among psyllid lineages. Genomes of several *Carsonella* lineages were revealed to be drastically reduced (158-174 kb), AT-rich (14.0–17.9% GC), and structurally conserved with similar gene inventories devoted for synthesizing essential amino acids scarce in the phloem sap. However, genomic information for secondary symbionts was limited. Hence, this study analyzed genomes of the bacteriome-associated dual symbionts, Secondary_AM (*Gammaproteobacteria*) and *Carsonella*_AM, in the mulberry psyllid *Anomoneura mori* (Psyllidae). This revealed that the Secondary_AM genome is as small and AT-rich (229,822 bp, 17.3% GC) as those of *Carsonella*, including *Carsonella*_AM (169,120 bp, 16.2% GC), implying that Secondary_AM is an evolutionarily ancient obligate mutualist like *Carsonella*. Phylogenomic analyses demonstrated that Secondary_AM is sister to “*Candidatus* Psyllophila symbiotica” of *Cacopsylla* spp. (Psyllidae), whose genomes (221–237 kb, 17.3–18.6% GC) were recently reported. The Secondary_AM and *Psyllophila* genomes showed highly conserved synteny, sharing all genes for complementing the incomplete tryptophan biosynthetic pathway of *Carsonella* and genes for synthesizing B vitamins. However, sulfur assimilation and carotenoid synthesizing genes were retained only in Secondary_AM and *Psyllophila*, respectively, indicating ongoing gene silencing. Average nucleotide identity, gene ortholog similarity, genome-wide synteny, and substitution rates suggested that the Secondary_AM/*Psyllophila* genomes are more labile than the *Carsonella* genomes.

## Introduction

Animals and microbes have diverse symbiotic relationships, among which the most intimate are bacteriome-associated mutualisms in insects (Douglas, 1989; Moran *et al*., 2008; Moran and Bennett, 2014; McCutcheon *et al*., 2019; Alarcón *et al*., 2022). Various insect lineages, especially those feeding on nutritionally restricted diets, such as plant sap and vertebrate blood, possess the bacteriome, a specialized organ with apparently varied developmental origins (Alarcón *et al*., 2022), which harbors the ‘primary symbiont’ providing nutritional supply to support the survival of holobionts (host-symbiont complexes). Distinct insect lineages harbor phylogenetically diverse primary symbionts, indicating their independent origins from various free-living microbes (Moran *et al*., 2008; Moran and Bennett, 2014; McCutcheon *et al*., 2019). They are mostly bacterial and feature organelle-like characteristics, including intracellular localization within the host, complete infection in host populations, host-symbiont cospeciation through strict vertical transmission, and drastic genome reduction due to elevated fixation of deleterious mutations and resulting loss of nonessential genes (Moran *et al*., 2008; Moran and Bennett, 2014; McCutcheon *et al*., 2019). Their highly reduced genomes retain many genes for synthesizing essential nutrients to fulfill the host requirements. However, the contingent acquisition of additional symbionts with a functionally intact genome may further allow the erosion of the primary symbiont genomes to the level where genes essential for the holobiont’s survival are also degraded and complemented by the newcomers. In that case, the primary symbionts *per se* may eventually be replaced by the newly acquired symbionts. Apparent snapshots of these evolutionary processes are observed in several insect lineages (McCutcheon *et al*., 2009; Vogel and Moran, 2013; Bennett *et al*., 2014; Koga and Moran, 2014; Husnik and McCutcheon, 2016; Bennett and Mao, 2018; Chong and Moran, 2018; Monnin *et al*., 2020; Michalik *et al*., 2021).

Psyllids (Hemiptera: Sternorrhyncha: Psylloidea) are phloem sap-sucking insects encompassing ∼4,000 described species worldwide (Burckhardt *et al*., 2021). They have a large bilobed bacteriome within the abdominal hemocoel (Profft, 1937; Buchner, 1965; Chang and Musgrave, 1969; Waku and Endo, 1987; Nakabachi *et al*., 2010; Dan *et al*., 2017; Nakabachi and Suzaki, 2023), which typically harbors two distinct bacterial symbionts (Subandiyah *et al*., 2000; Spaulding and von Dohlen, 2001; Hall *et al*., 2016; Morrow *et al*., 2017; Nakabachi *et al*., 2022a; Dittmer *et al*., 2023; Maruyama *et al*., 2023). The primary symbiont is “*Candidatus* Carsonella ruddii” (*Gammaproteobacteria*: *Oceanospirillales*, hereafter *Carsonella*) (Thao, Moran, *et al*., 2000), which has been detected in all psyllid species analyzed to date and is thus thought essential for Psylloidea (Spaulding and von Dohlen, 2001; Sloan and Moran, 2012b; Nakabachi, Ueoka, *et al*., 2013; Arp *et al*., 2014; Hall *et al*., 2016; Morrow *et al*., 2017; Nakabachi and Moran, 2022; Nakabachi *et al*., 2022b, 2022a). Molecular phylogenetic analyses demonstrated cospeciation between *Carsonella* lineages and their host psyllids, resulting from a single acquisition of an ancestor of *Carsonella* by a psyllid common ancestor and its subsequent strict vertical transmission (Thao, Moran, *et al*., 2000; Spaulding and von Dohlen, 2001; Hall *et al*., 2016; Morrow *et al*., 2017; Maruyama *et al*., 2023). The *Carsonella* genomes derived from several psyllid lineages were analyzed and revealed to be drastically reduced (158–174 kb), AT-rich (14.0–17.9% GC), and structurally conserved. They lack numerous genes apparently essential for bacterial life but retain genes for synthesizing essential amino acids that are deficient in the phloem sap diet (Nakabachi *et al*., 2006; Nakabachi, 2008; Sloan and Moran, 2012b; Nakabachi, Ueoka, *et al*., 2013; Nakabachi, Piel, *et al*., 2020; Dittmer *et al*., 2023). Another bacterial lineage housed in the bacteriome is categorized as a ‘secondary symbiont,’ which is phylogenetically diverse among psyllid lineages, suggesting its repeated infections and replacements during the evolution of Psylloidea (Thao, Clark, *et al*., 2000; Spaulding and von Dohlen, 2001; Sloan and Moran, 2012b; Hall *et al*., 2016; Morrow *et al*., 2017; Nakabachi, Malenovský, *et al*., 2020; Nakabachi *et al*., 2022b, 2022a). Although secondary symbionts in diverse insect taxa form various host-symbiont relationships across the mutualism-parasitism continuum (Gherna *et al*., 1991; Dale and Maudlin, 1999; Nakabachi *et al*., 2003; M. L. L. Thao and Baumann, 2004; Moran *et al*., 2008; Werren *et al*., 2008; Oliver *et al*., 2010; Johnson, 2015), those in the psyllid bacteriome appear to be consistently hosted by all individuals within a particular psyllid species, having obligate mutualistic relationships with the host psyllid (Sloan and Moran, 2012b; Nakabachi, Nikoh, *et al*., 2013; Nakabachi, Ueoka, *et al*., 2013; Nakabachi, Piel, *et al*., 2020; Dittmer *et al*., 2023). Therefore, they are occasionally called ‘co-primary symbionts,’ as in various auchenorrhynchan insects (McCutcheon *et al*., 2009; Bennett *et al*., 2014; Bennett and Mao, 2018; Michalik *et al*., 2021). Contrasting to *Carsonella*, genomic information of secondary symbionts associated with psyllid bacteriomes was limited. Sloan and Moran determined the whole genome sequences of two distantly related secondary symbionts (both *Gammaproteobacteria*: *Enterobacterales*) derived from *Ctenarytaina eucalypti* (Aphalaridae: Spondyliaspidinae) and *Heteropsylla cubana* (Psyllidae: Ciriacreminae) (Sloan and Moran, 2012b). This revealed that they are relatively large with less biased nucleotide contents (1441 kb with 43.3% GC in *C. eucalypti*; 1122 kb with 28.9% GC in *H. cubana*) compared to other insect bacteriome-associated symbionts analyzed thus far, including *Carsonella*, suggesting their recently formed relationships with the host psyllid. Both genomes encoded genes to complement the incomplete essential amino acid biosynthetic pathways encoded in the genome of co-residing *Carsonella* (Sloan and Moran, 2012b). Subsequently, analyses of two lineages of “*Candidatus* Profftella armatura” (*Gammaproteobacteria*: *Burkholderiales*, hereafter *Profftella*) from the Asian citrus psyllid *Diaphorina citri* and its relative *Diaphorina* cf. *continua* (both Psyllidae: Diaphorininae) revealed that their genomes are highly reduced and AT-rich (460 kb with 24.2% GC in *D. citri*; 470 kb with 24.4% GC in *D*. cf. *continua*) (Nakabachi, Ueoka, *et al*., 2013; Nakabachi, Piel, *et al*., 2020), suggesting that *Profftella* is an ancient symbiont. The highly conserved synteny and comparatively low substitution rates of the genomes implied that *Profftella* has acquired a relatively stable status, which is comparable to that of *Carsonella*. These genomes shared all genes for the biosynthesis of toxins (diaphorin and hemolysin), carotenoids, and B vitamins (riboflavin and biotin), indicating that *Profftella* is a unique versatile symbiont playing multiple roles, including nutritional supply and defense against natural enemies. Subsequent studies have demonstrated distinct activities of diaphorin, including inhibitory effects against divergent organisms (Nakabachi and Fujikami, 2019; Nakabachi and Okamura, 2019; Yamada *et al*., 2019; Tanabe *et al*., 2022; Takasu *et al*., 2023, 2024). These collective benefits to the host may contribute to stabilizing symbiotic relationships, leading to the organelle-like status of *Profftella* (Nakabachi, Ueoka, *et al*., 2013; Nakabachi, Piel, *et al*., 2020). However, these four lineages reported previously were only a tiny fraction of the diverse secondary symbionts associated with the psyllid bacteriome, making it uncertain about their evolutionary behaviors and how common versatile symbionts like *Profftella* are in Psylloidea.

Thus, as a first step to obtaining more insights into the evolution of the dual symbiotic system of the psyllid bacteriome, we analyzed the genomes of the bacteriome-associated secondary symbiont (*Gammaproteobacteria*: *Enterobacterales*, hereafter Secondary_AM) and *Carsonella* (hereafter *Carsonella*_AM) of a sericultural pest, the mulberry psyllid *Anomoneura mori* (Psyllidae: Psyllinae), in which the localization of the symbionts was determined (Fukatsu and Nikoh, 1998). During the analysis, genomes of 12 *Carsonella* lineages and 10 lineages of a secondary symbiont “*Candidatus* Psyllophila symbiotica” (*Gammaproteobacteria*: *Enterobacterales*, hereafter *Psyllophila*) from four *Cacopsylla* spp. (Psyllidae: Psyllinae) were recently published, revealing that the *Psyllophila* genomes are highly reduced (221–237 kb) and AT-rich (17.3–18.6% GC), encoding genes complementary to those of *Carsonella* (Dittmer *et al*., 2023). We compared the genomes of the bacteriome-associated symbionts of *A. mori* with these previously sequenced genomes of psyllid symbionts.

## Materials and Methods

### Insect specimen and DNA preparation

The material of *Anomoneura mori* was collected from the mulberry tree *Morus* sp. (Moraceae) in Tsukuba City, Ibaraki Prefecture, Honshu, Japan (36.048N, 140.101E, 23 m a.s.l.) on May 26, 2015. DNA was extracted from a pool of bacteriomes of three adult female *A. mori* using a DNeasy Blood & Tissue Kit (Qiagen) following the manufacturer’s instructions. Whole genome amplification was performed using the extracted DNA and a REPLI-g Mini Kit (Qiagen) according to the manufacturer’s instructions.

### Sequencing and assembly

An 800-bp paired-end library and an 8-kbp mate-pair library of the *A. mori* bacteriome DNA were prepared using the TruSeq DNA PCR-Free Sample Preparation kit (Illumina) and Nextera Mate Pair Sample Preparation kit (Illumina), respectively. The libraries were sequenced on the MiSeq instrument (Illumina) with the MiSeq Reagent kit v3 (600-cycles; Illumina). From obtained sequences, paired reads in which either of the pair showed similarity (e-value scores < 1.0E-5) to previously reported genomic sequences of eight *Carsonella* lineages (NC_018417.1, AP009180.1, CP003541.1, CP003542.1, CP003543.1, CP003545.1, CP003467.1, CP012411.1) were collected using local alignment search with Nucleotide Basic Local Alignment Search Tool (BLASTN) program (Camacho *et al*., 2009) with a custom Perl script. Sequencing errors were corrected using ShortReadManager based on 17-mer frequency (Ohtsubo *et al*., 2022). These collected and refined reads were assembled using Newbler version 2.9 (Roche). Gap sequences between contigs were determined *in silico* using GenoFinisher and AceFileViewer (Ohtsubo *et al*., 2022).

### Annotation and structural analysis of the genomes

Initial gene predictions and annotations were conducted using the DNA Data Bank of Japan (DDBJ) Fast Annotation and Submission Tool (DFAST) pipeline version 1.2.0 (Tanizawa *et al*., 2018) and the National Center for Biotechnology Information (NCBI) Prokaryotic Genome Annotation Pipeline (PGAP) version 2023-05-17.build6771 (Tatusova *et al*., 2016), followed by manual corrections with the aid of Rfam version 12.2 (Kalvari *et al*., 2021), the NCBI ORFfinder (Wheeler *et al*., 2003), BLAST (Camacho *et al*., 2009), and eggNOG-mapper version 2.1.12 (Cantalapiedra *et al*., 2021). Functional categories of Cluster of Orthologous Groups (COG) were assigned to predicted genes using the abovementioned version of eggNOG-mapper (Cantalapiedra *et al*., 2021). Metabolic pathways were analyzed using the Kyoto Encyclopedia of Genes and Genomes (KEGG) (Kanehisa, 2019). Pathway maps were created by examining the presence or absence of genes involved in the biosynthesis of essential amino acids and vitamins. Dinucleotide bias and GC skew were analyzed and circular diagrams were drawn using ArcWithColor version 1.62 (Ohtsubo *et al*., 2008). The codon adaptation index (CAI) was calculated using the CAIcal server (Puigbo *et al*., 2008). Pairwise comparisons of genomic structures were performed using GenomeMatcher version 3.06 (Ohtsubo *et al*., 2008), in which BLASTN of all-against-all bl2seq similarity searches were conducted with the parameter set ‘-F F -W 21 -e 1.0e-10’.

### Phylogenomic analysis

Single-copy orthologous proteins shared among Secondary_AM, ten lineages of *Psyllophila* from four *Cacopsylla* spp. (six from *C. melanoneura*, two from *C. picta*, and a single each from *C. pyricola* and *C. pyri*) (Dittmer *et al*., 2023), 33 other insect nutritional endosymbionts belonging to the order *Enterobacterales*, and *Pseudomonas oryziphila* and two strains of *Pseudomonas entomophila* (both *Gammaproteobacteria*: *Pseudomonadales*) as outgroups were identified using OrthoFinder version 2.5.5 (Emms and Kelly, 2019). The amino acid sequences of identified proteins were aligned with MAFFT 7.452 (Katoh and Standley, 2013) using the E-INS-i algorithm with the aid of Perl script MultiMafft.pl (https://fish-evol.org/MAFFT.html). After removing amino acid sites corresponding to alignment gaps, the multiple fasta alignments were concatenated into a partitioned supermatrix using catsequences version 1.4 (https://github.com/ChrisCreevey/catsequences.git). The Phylogenetic trees were inferred by the Maximum Likelihood (ML) method using RAxML-NG version 1.2.1 (Kozlov *et al*., 2019) with 1,000 replicates with the MTREV+I+G4+F model, which was selected using ModelTest-NG version 0.1.7 (Darriba *et al*., 2020). Trees were visualized using Interactive Tree of Life (iTOL) version 5 (Letunic and Bork, 2021).

### Average nucleotide identity analysis

To measure the nucleotide-level genomic similarity of *A. mori* symbionts to their counterparts in four *Cacopsylla* spp. (ten *Carsonella* lineages as mentioned above and eight *Psyllophila* lineages from *C. melanoneura*, two from *C. picta*, and a single each from *C. pyricola* and *C. pyri*), the average nucleotide identity (ANI) was calculated using the ANI calculator (Rodriguez-R and Konstantinidis, 2016). The calculation was performed using reciprocal best hits (two-way ANI) between two genomic datasets with the default parameters. Window and step sizes were set to 1000 bp and 200 bp, respectively.

### Substitution rate analysis

Amino acid sequences were deduced from the protein-coding genes (CDSs) shared between the *A. mori* symbionts and their counterparts in the melET isolate of *C. melanoneura*, which were then aligned with MAFFT version 7.452 (Katoh and Standley, 2013) using the E-INS-i algorithm with default parameters. The resulting protein alignments were converted to nucleotide alignments using PAL2NAL version 13.0 (Suyama *et al*., 2006). Nonsynonymous (*dN*) and synonymous (*dS*) substitution rates and *dN*/*dS* ratios between orthologous pairs were calculated using the KaKs_Calculator 1.2 package (Zhang *et al*., 2006) with the YN model (Yang and Nielsen, 2000). All statistical analyses were performed using the R software version 4.2.1 (R Core Team 2022, https://www.r-project.org).

## Results

### The Secondary_AM genome is as extremely small and AT-rich as the *Carsonella* genomes

MiSeq sequencing of the *A. mori* bacteriome DNA libraries yielded 1.36 million paired-end reads Mbp) and 1.03 million mate-pair reads (447 Mbp). From these, we collected sequence reads that showed detectable similarity (BLASTN e-value scores < 1.0E-5) to published *Carsonella* genomes, aiming to determine the *Carsonella*_AM genome. The assemblage of obtained and refined 126 thousand paired-end reads (72.6 Mbp) and 25 thousand mate pair reads (5.58 Mbp) yielded five scaffolds and 220 large contigs (> 0.5 kbp). After filling gap sequences *in silico*, somewhat unexpectedly, we succeeded in obtaining circular complete genomes with the coverage of ca. 100× not only *Carsonella*_AM (169,120 bp, 16.2% GC) but also Secondary_AM (229,822 bp, 17.3% GC) (Table 1, Fig 1), which were identified by the presence of known sequences of 16S rRNA genes (Fukatsu and Nikoh, 1998; Nakabachi *et al*., 2022b). The *Carsonella*_AM genome encoded 196 predicted protein-coding sequences (CDSs), a single rRNA operon, and 28 tRNAs (Tables 1 and S1, Fig 1). The Secondary_AM genome encoded 215 CDSs, a single rRNA operon, and 26 tRNAs (Tables 1 and S2, Fig 1). Notably, the size and GC content of the Secondary_AM genome were comparable to those of *Carsonella*, including *Carsonella*_AM, implying an evolutionarily ancient and obligate mutualistic relationship of Secondary_AM with the host insect. These features were similar to those of recently published genomes of *Psyllophila* (221,413−237,114 bp, 17.3−18.6% GC), the bacteriome-associated secondary symbiont of *Cacopsylla* spp. (Psyllidae: Psyllinae) (Dittmer *et al*., 2023), starkly contrasting to the previously sequenced much larger genomes with less nucleotide composition bias, derived from secondary symbionts of two psyllid species, *Ctenarytaina eucalypti* (Aphalaridae: Spondyliaspidinae) (1.4 Mb, 43.3% GC) and *Heteropsylla cubana* (Psyllidae: Ciriacreminae) (1.1 Mb, 28.9% GC) (Sloan and Moran, 2012b).

**Fig. 1.-.**
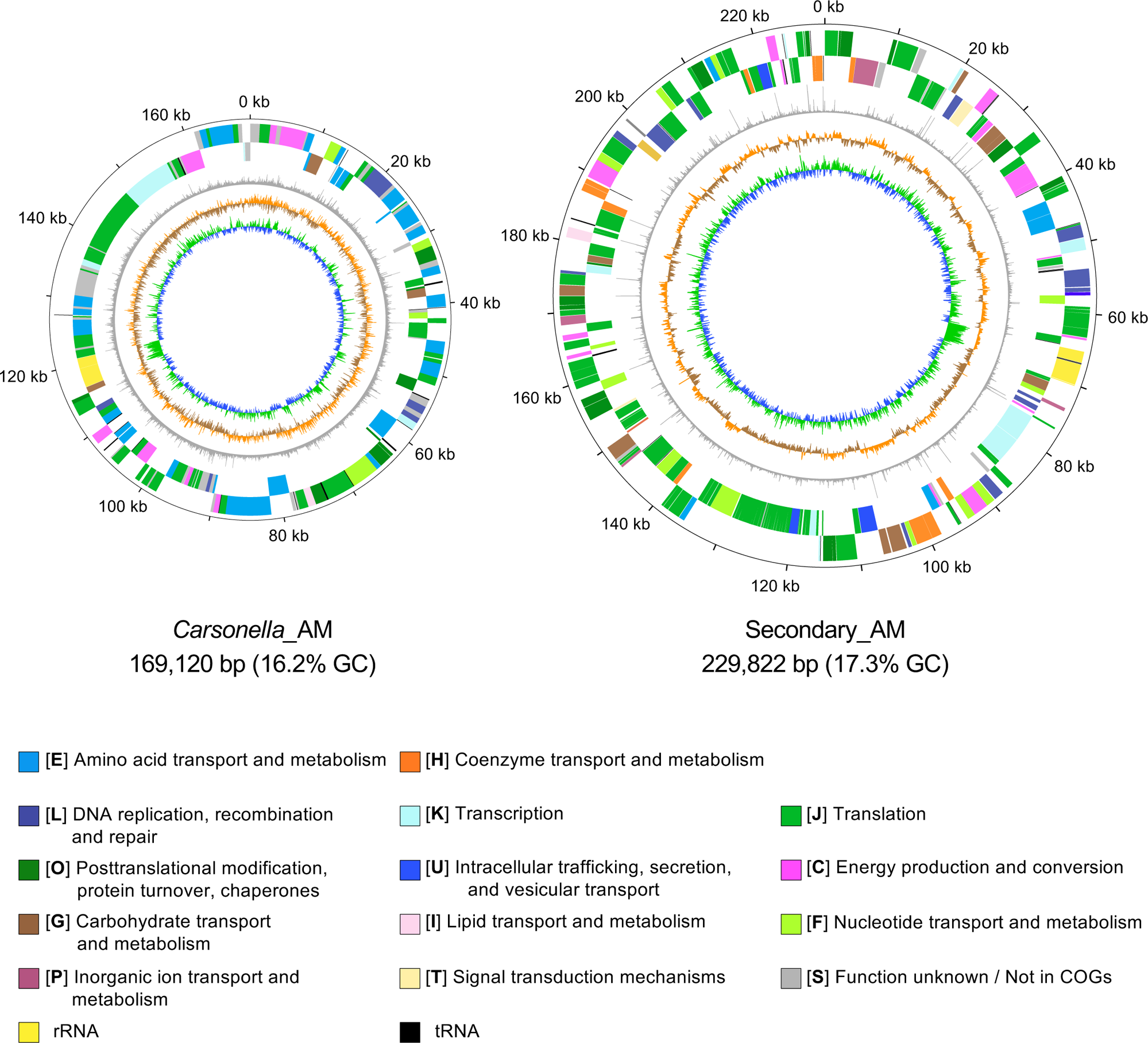
Circular representation of the genomes of Secondary_AM and *Carsonella*_AM. The concentric rings denote the following features (from the outside): (i) the scale in kilobases, (ii) forward strand genes, (iii) reverse strand genes, (iv) dinucleotide bias, (v) GC skew, and (vi) G + C content. For the calculation of (iv), (v), and (vi), sliding windows of 100 bp and a step size of 10 bp were used.

**Table 1.** Genomic features of the bacteriome-associated symbionts in *A. mori*.

|  | *Carsonella*_AM | Secondary_AM |
| --- | --- | --- |
| Chromosome (bp) | 169,120 | 229,822 |
| G+C content (%) | 16.2 | 17.3 |
| CDS | 196 | 215 |
| rRNA | 3 | 3 |
| tRNA | 28 | 26 |

### Secondary_AM is sister to *Psyllophila*

As previous phylogenetic analyses using 16S rRNA genes showed relatedness between Secondary_AM and *Psyllophila* (Nakabachi *et al*., 2022b; Schuler *et al*., 2022; Štarhová Serbina *et al*., 2022), we performed more detailed phylogenetic analysis using sequences for orthologous proteins encoded in these genomes. OrthoFinder v2.5.5 (Emms and Kelly, 2019) identified 60 single-copy orthologous proteins shared among Secondary_AM, 10 *Psyllophila* lineages from four *Cacopsylla* spp. (Dittmer *et al*., 2023), 33 other *Enterobacterales* endosymbionts of insects, and *P. oryziphila* and two strains of *P. entomophila* (all *Pseudomonadales*) (Table S2). After aligning these single-copy orthologous proteins and trimming poorly aligned regions, a total of 14,766 amino acid positions were concatenated, on which a maximum likelihood phylogenetic analysis was performed (Fig 2). The results showed that Secondary_AM forms a robustly supported clade (bootstrap: 100%) with a clade formed by 10 *Psyllophila* lineages derived from *Cacopsylla* spp., which is also robustly supported (bootstrap: 100%) in the ML tree. This branching pattern indicates that Secondary_AM is a sister lineage of *Psyllophila*, sharing a recent common ancestor. The clade of Secondary_AM and *Psyllophila* further formed a robustly supported clade (bootstrap: 100%) with “*Candidatus* Annandia” spp., the bacteriome-associated symbionts of adelgids (Hemiptera: Sternorrhyncha: Phylloxeroidea: Adelgidae), and “*Candidatus* Nardonella” spp., the bacteriome-associated symbionts of weevils (Coleoptera: Curculionoidea). The previously sequenced secondary symbionts of *C. eucalypti* and *H. cubana* were shown to be distantly related to this clade encompassing Secondary_AM and *Psyllophila* (Fig 2).

**Fig. 2.-.**
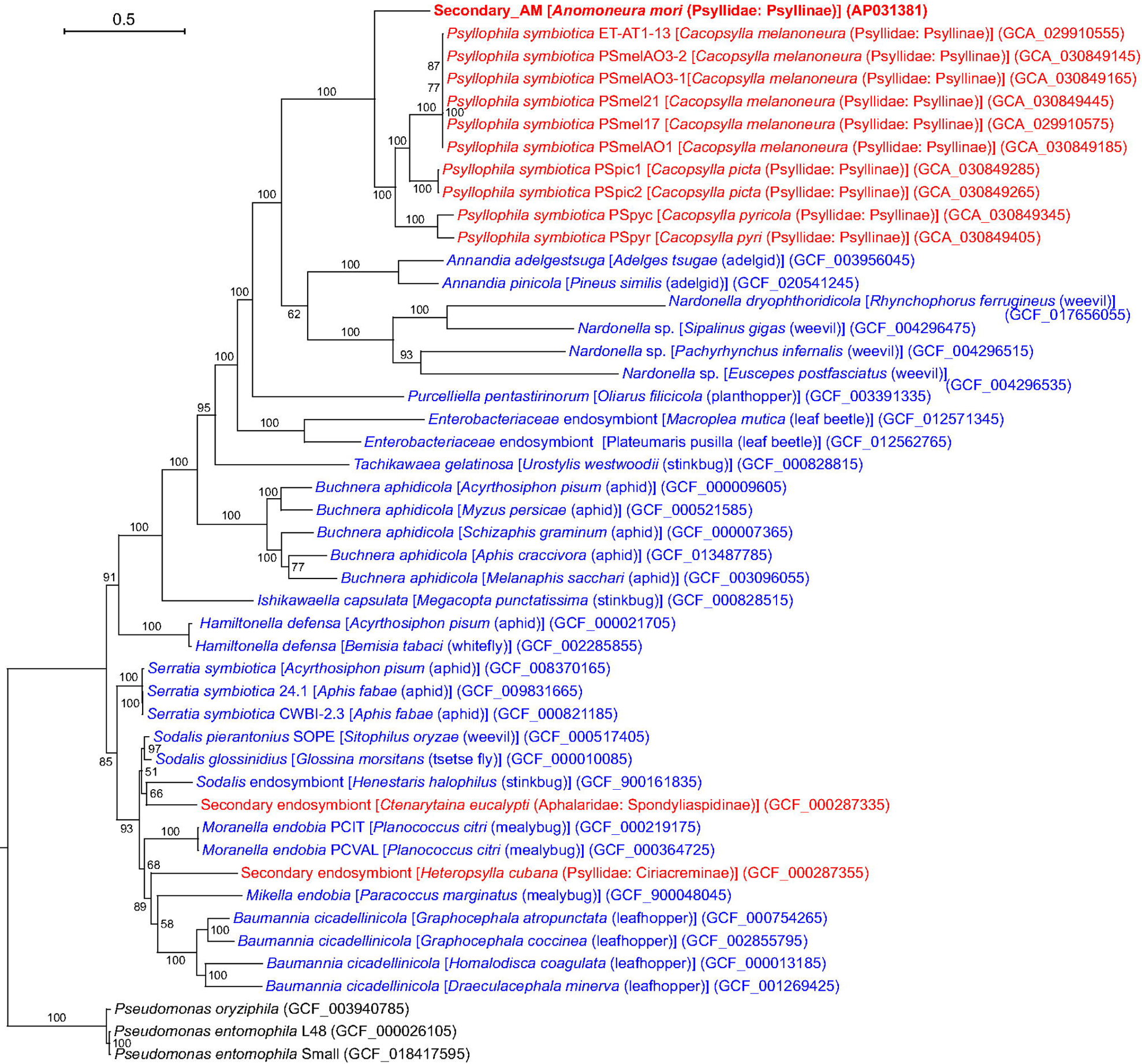
Maximum likelihood phylogram showing phylogenetic relationships among Secondary_AM, 10 *Psyllophila* lineages from four *Cacopsylla* spp., and 33 other insect endosymbionts belonging to the order Enterobacterales. For simplicity, species names are presented without “*Candidatus*.” A total of 14,766 unambiguously aligned amino acid positions derived from 60 single-copy orthologous proteins shared among these bacteria were subjected to the analysis. On each branch, bootstrap support values are shown. The scale bar indicates substitutions per site. Regarding symbiotic bacteria, host organisms are shown in brackets. Symbionts of animals other than psyllids are shown in blue, while symbionts of psyllids are shown in red. Sequences from the present study are shown in bold. DDBJ/EMBL/GenBank accession numbers are provided in parentheses. *Pseudomonas oryziphila* and two strains of *Pseudomonas entomophila* (all Gammaproteobacteria: Pseudomonadales) were used as an outgroup.

To further assess the genomic similarity between Secondary_AM and 10 *Psyllophila* lineages, the average nucleotide identity (ANI) (Jain *et al*., 2018) was calculated (Table 2). For comparison, ANI was calculated also between *Carsonella*_AM and 12 *Carsonella* lineages from *Cacopsylla* spp. The results showed that 1) the mean ANI is higher in *Carsonella* (86.915 ± 0.005%) [mean ± standard deviation (SD); *n* = 12] than in *Psyllophila* (79.571 ± 0.004%; *n* = 10) (*p* < 0.001, Welch’s *t*-test), and 2) the ANI values are close to one another within *Carsonella* or *Psyllophila* (see SDs of 1 and Table 2). However, as the ANI of Secondary_AM was highest (79.78%) with the *Psyllophila* lineage derived from the melET isolate of *C. melanoneura* (PSmelET) (Table 2), we further compared their sequences in more detail, featuring this lineage.

**Table 2.**
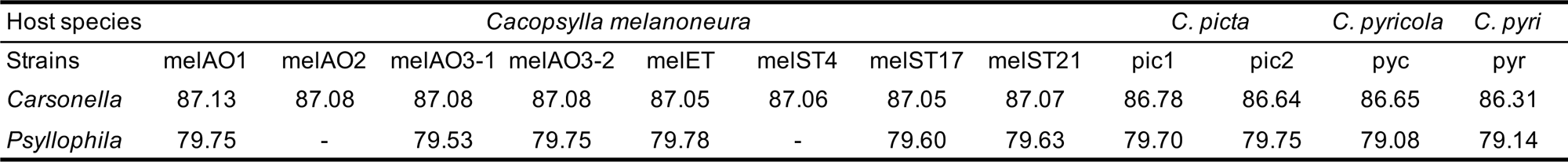
Average nucleotide identity between symbionts of *A. mori* and *Cacopsylla* spp.

### Secondary_AM and *Psyllophila* exhibit conserved genomic synteny

Genome-wide alignments revealed the highly conserved synteny between the genomes of Secondary_AM and 10 *Psyllophila* lineages (the alignment with the PSmelET genome is shown as a representative in Fig 3), indicating that most genes are shared between these symbionts, and essentially no genome rearrangements have occurred since they diverged. Between the genomes of Secondary_AM and PSmelET, a total of 230 pairs of orthologous genes were 79.8% identical on average at the nucleotide level, and amino acid sequences of 200 pairs of orthologous proteins were 69.0% identical on average (Table S2). They shared all genes involved in synthesizing biotin and riboflavin or complementing the essential amino acid biosynthesis by *Carsonella*, which will be further discussed later.

**Fig. 3.-.**
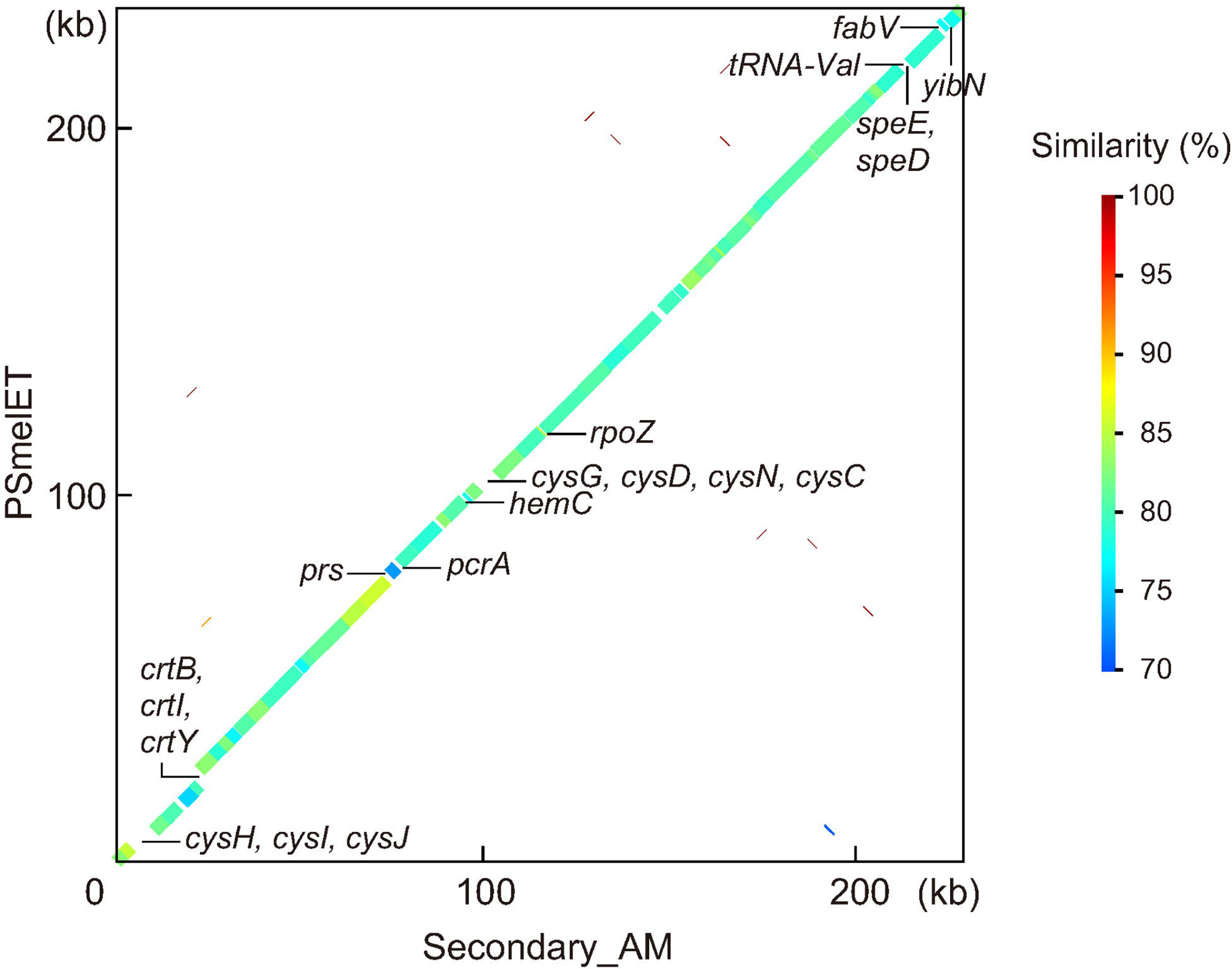
Comparison of the genomic structures of Secondary_AM and *Psyllophila* derived from *Cacopsylla melanoneura* (PSmelET). The genomes of Secondary_AM and PSmelET are represented by the x and y axes, respectively. The thick line indicates the shared synteny between the two genomes. The color of the line indicates the percentage similarity between the nucleotide sequences. The genes found in Secondary_AM, but not in PSmelET, are presented below the line plot; the genes present in PSmelET, but not in Secondary_AM, are shown above the line plot.

Despite this high level of conservation between the genomes, random gene silencing appeared ongoing in Secondary_AM and *Psyllophila*. The genes found in one of the lineages but not in the other are shown in Fig 3. These genes showed no signs of horizontal acquisition after the divergence of these symbionts, indicating that the different gene inventories reflect gene silencing on either genome. Notably, *cysCDHIJN* encoding enzymes for sulfur assimilation were retained in Secondary_AM but were missing in PSmelET and all other *Psyllophila* lineages (Figs 3 and S2, Table S2). These genes potentially contribute to synthesizing sulfur-containing amino acids, cysteine and methionine, the latter of which is an essential amino acid that the host insects cannot synthesize. However, no other genes related to their synthesis were retained in the Secondary_AM genome (Fig S2, Table S2). Moreover, *Carsonella*_AM lacked genes for the biosynthesis of cysteine/methionine other than *metE*, which converts homocysteine into methionine (Figs 4 and S2). Thus, the cysteine/methionine synthesis pathway appeared incomplete, making the role of the conservation of *cysCDHIJN* genes in Secondary_AM unclear.

**Fig. 4.-.**
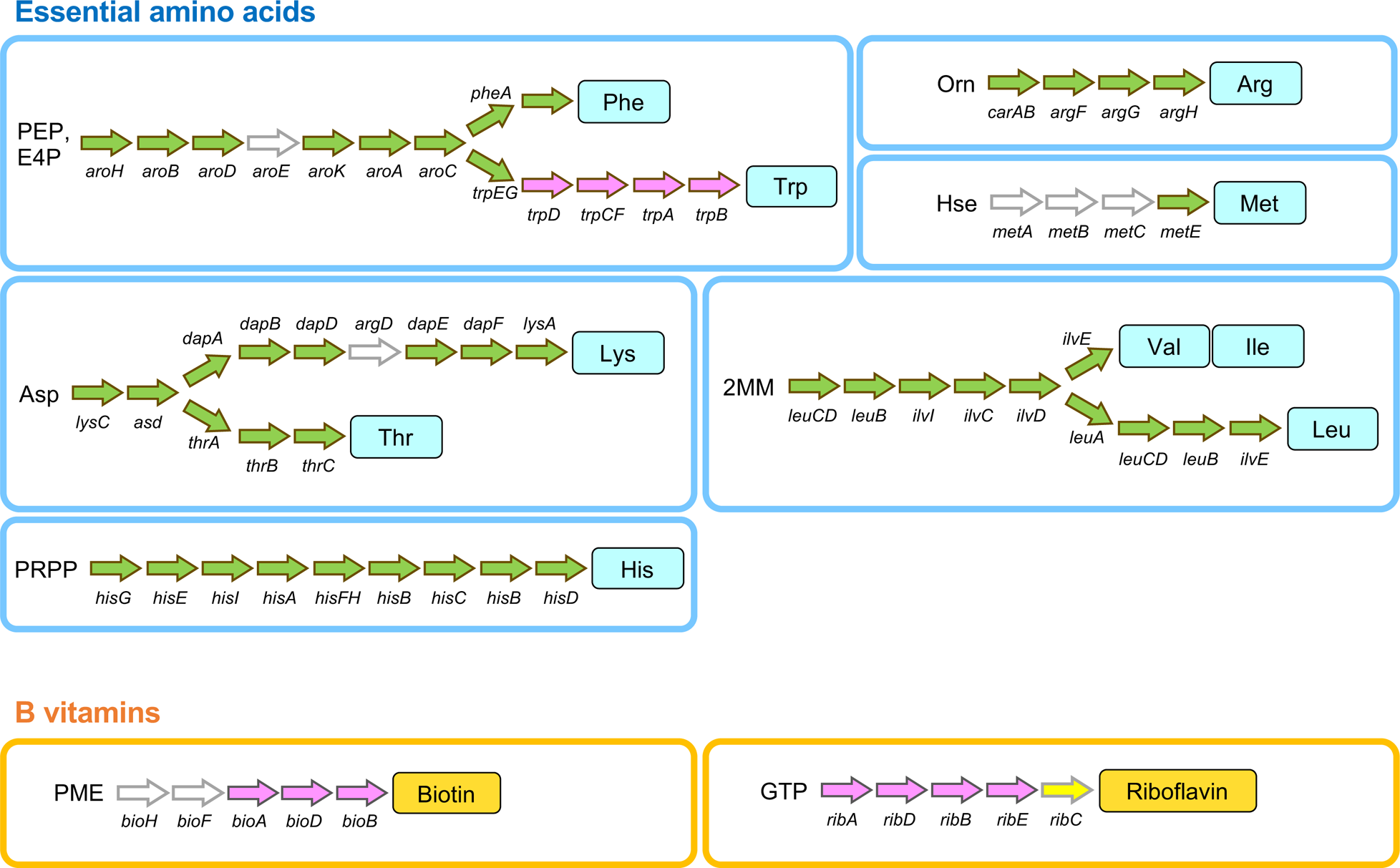
Pathways for synthesizing essential amino acids and B vitamins reconstructed from genes encoded in the genomes of *Carsonella*_AM and Secondary_AM. Genes denoted with green and magenta arrows are found in the genomes of *Carsonella*_AM and Secondary_AM, respectively. Those shown with white arrows appear to be absent in these genomes. The *ribC* shown with a yellow arrow is suspected to be encoded in the host psyllid genome, which was horizontally acquired from an unknown bacterium. PEP, phosphoenolpyruvate; E4P, erythrose 4-phosphate; 2MM, 2-methylmalate; PRPP, phosphoribosyl diphosphate; PME, pimelyl-(acyl-carrier protein) methyl ester.

On the other hand, the genomes of all *Psyllophila* lineages, including PSmelET, retained genes for synthesizing carotenoids (*crtB*, *crtI*, and *crtY*), all of which were missing from Secondary_AM (Fig 3), indicating that the lineage leading to Secondary_AM has lost the ability to synthesize carotenoids.

### *Carsonella*_AM and Secondary_AM are metabolically interdependent

Among the COG functional categories assigned to protein-coding genes of *Carsonella*_AM and Secondary_AM, “translation” (category J) exhibited the highest percentage in both symbionts (33.2% in *Carsonella*_AM, 45.5% in Secondary_AM), while those for “transcription” (category K, 2.5% in *Carsonella*_AM, 3.6% in Secondary_AM) and DNA replication, recombination and repair (category L, 2.5% in *Carsonella*_AM, 6.4% in Secondary_AM) were relatively low (Fig S1). This implies that translation is an especially important information-processing process for retaining the autonomy of symbionts within the host cell. Indeed, high levels of conservation of translation genes have been observed in diverse heritable bacterial symbionts of insects (Nakabachi *et al*., 2006; Toft *et al*., 2009; McCutcheon and Moran, 2012; Rao *et al*., 2015; Santos-Garcia *et al*., 2015; Weglarz *et al*., 2018), contrasting the case of free-living bacteria in which the percentages for these COG categories regarding information processing are comparable and ∼5% (Tatusov *et al*., 1997, 2000; Nakabachi, 2008; Wright *et al*., 2021; Thiruvengadam *et al*., 2022). In *Carsonella*_AM, the second most prominently represented COG category was E “amino acid transport and metabolism” (21.3%) (Fig S1). As with other *Carsonella* strains previously analyzed (Nakabachi *et al*., 2006; Nakabachi, 2008; Sloan and Moran, 2012b; Nakabachi, Ueoka, *et al*., 2013; Nakabachi, Piel, *et al*., 2020; Dittmer *et al*., 2023), *Carsonella*_AM retained most genes required for the biosynthesis of essential amino acids, although some appeared missing (Figs 4 and S2, Table S1). Regarding the tryptophan biosynthesis pathway, only *trpE* and *trpG*, genes for anthranilate synthase components involved in the first catalytic step after diverging from that for phenylalanine, were retained in the *Carsonella*_AM genome. All other genes (*trpD, trpCF, trpA, and trpB*) required for the rest of the pathway were missing. Notably, these genes lost in *Carsonella*_AM were retained entirely in the Secondary_AM genome (Figs 4 and S2). Moreover, these were the only Secondary_AM genes directly involved in synthesizing essential amino acids, underlining the elaborate metabolic complementarity encoded in the genomes of *Carsonella*_AM and Secondary_AM. Interestingly, the interdependent biosynthetic pathway of tryptophan, involving *Carsonella* encoding only *trpE* and *trpG*, complemented by a secondary symbiont, was observed not only in *Cacopsylla* spp. but also in *C. eucalypti* (Aphalaridae: Spondyliaspidinae) and *H. cubana* (Psyllidae: Ciriacreminae) with distantly related and possibly more recently acquired *Enterobacterales* symbionts with larger genomes (Sloan and Moran, 2012b), presenting notable examples of convergence.

Although only 5.5% of genes encoded in the Secondary_AM genome were assigned with the COG category H “coenzyme transport and metabolism,” they were *bioA*, *bioD*, and *bioB*, which are required for synthesizing a B vitamin, biotin, and *ribA*, *ribD*, *ribB*, and *ribE*, genes for the synthesis of another B vitamin, riboflavin, suggesting that the synthesis of B vitamins, which are also scarce in the phloem sap diet (Ziegler *et al*., 1975), is the pivotal role of Secondary_AM. All of these genes were retained not only in *Psyllophila* but also in *Profftella*, a distantly related symbiont whose primary role appears to be the protection of the holobiont from natural enemies, further adding an example of intriguing convergence (Figs 4 and S2). Although all the lineages of Secondary_AM, *Psyllophila*, and *Profftella* lacked *ribC* that is required for the final step of riboflavin biosynthesis, a previous screening of genomic/transcriptomic data of several divergent psyllid lineages demonstrated that host psyllids horizontally acquired *ribC* from an uncertain bacterial lineage before the radiation of the major psyllid lineages (Sloan *et al*., 2014; Nakabachi, 2015), which is supposed to complement its absence in bacterial symbionts.

### The Secondary_AM/*Psyllophila* genomes are more labile than *Carsonella*

Although diverse lineages of insect primary symbionts have been demonstrated to have highly stable genomic structures (Tamas et al. 2002; Degnan et al. 2005; Rio et al. 2012; Sloan & Moran 2012b; Bennett & Moran 2013; Nakabachi, Ueoka, et al. 2013; Koga & Moran 2014; Husnik & McCutcheon 2016; Mao et al. 2017), this is usually not the case for more recently acquired secondary symbionts (Burke and Moran, 2011; Manzano-marin *et al*., 2012; Bennett *et al*., 2014; Koga and Moran, 2014; Campbell *et al*., 2015; McCutcheon *et al*., 2019; Nakabachi, Piel, *et al*., 2020). When compared between symbionts of *A. mori* and their counterparts in the melET isolate of *C. melanoneura*, for which the same divergence time is assumed to be applicable, ANI (Table 2), genome-wide synteny (Figs 3 and S3), and similarity of orthologous genes [87.3 ± 9.9% (mean ± SD; *n* = 215) for *Carsonella* vs. 79.8 ± 9.5% for Secondary_AM/*Psyllophila* (*n* = 230)] (Tables S1 and S2) suggested that the *Carsonella* genomes are more stable than the Secondary_AM/*Psyllophila* genomes. To further assess the genomic stability, we analyzed the genome-wide rates of synonymous (*dS*) and nonsynonymous (*dN*) substitutions for these symbionts (Fig 5, Tables S1 and S2). Orthologous protein-coding genes, namely, 185 pairs of *Carsonella* genes and 200 pairs of the Secondary_AM/*Psyllophila* genes, were used for the analysis. The mean values of both *dN* (0.076 ± 0.046 for *Carsonella* vs. 0.148 ± 0.053 for Secondary_AM/*Psyllophila*) and *dS* (2.84 ± 12.34 for *Carsonella* vs. 13.16 ± 30.36 for Secondary_AM/*Psyllophila*) were higher (*p* < 0.001, Brunner-Munzel test) in the Secondary_AM/*Psyllophila* lineage than in *Carsonella*. Synonymous divergence appeared saturated (*dS* > 3.0) in 15 (8.1%) *Carsonella* genes and 95 (47.5%) Secondary_AM/*Psyllophila* genes (Tables S1 and S2). To be conservative, only genes with *dS* < 3.0 were used for calculating *dN*/*dS*, which was also higher (*p* < 0.001, Brunner-Munzel test) in the Secondary_AM/*Psyllophila* lineages (0.134 ± 0.111, *n* = 105) than in *Carsonella* (0.121 ± 0.151, *n* = 170) (Fig 5, Tables S1 and S2). However, no gene was estimated to have *dN*/*dS* > 1, indicating purifying selection for all genes analyzed in these symbionts.

**Fig. 5.-.**
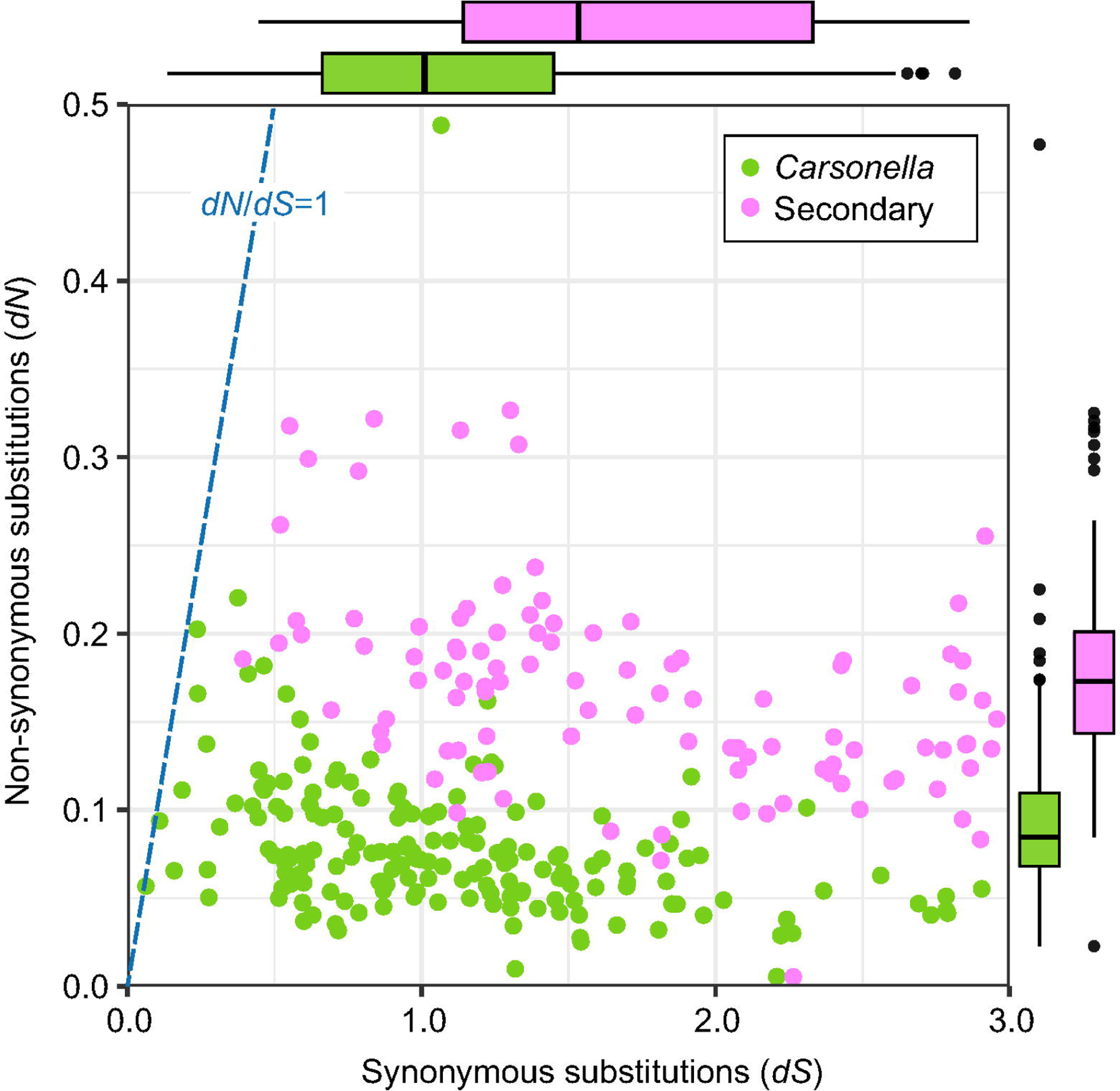
Synonymous (*dS*) and nonsynonymous (*dN*) substitution rates inferred from pairwise comparisons of orthologous CDSs in the Secondary_AM/ *Psyllophila* lineages (magenta dots) and *Carsonella* (green dots). Values of genes with *dS* < 3.0 (104 and 172 orthologous pairs in Secondary_AM/*Psyllophila* and *Carsonella*, respectively), which were used for calculating *dN*/*dS*, are shown. Box plots (Secondary_AM/*Psyllophila*, magenta; *Carsonella*, green) on the x- and y-axes indicate distributions (median, quartiles, minimum, maximum, and outliers) of *dS* and *dN* values, respectively.

## Discussion

The present study determined the complete genome sequences of Secondary_AM and *Carsonella*_AM, the symbionts harbored in the *A. mori* bacteriome, which revealed that the Secondary_AM genome is as small and AT-rich (229,822 bp, 17.3% GC) as those of *Carsonella* lineages (158–174 kb, 14.0–17.9% GC), including *Carsonella*_AM (169,120 bp, 16.2% GC) (Table 1, Fig 1), implying that Secondary_AM is also an evolutionarily ancient obligate mutualist for the host insect. The molecular phylogenetic analysis using 60 single-copy orthologous proteins demonstrated that Secondary_AM is sister to *Psyllophila* lineages derived from *Cacopsylla* spp. (Fig 2). Previous microbiome and phylogenetic analyses using 16S rRNA genes suggested that the Secondary_AM/Psyllophila lineage is widely distributed in *Cacopsylla* spp. (Nakabachi *et al*., 2022b; Schuler *et al*., 2022; Štarhová Serbina *et al*., 2022) and also in some species of other genera, including *Anomoneura* and *Cyamophila* (all Psyllidae: Psyllinae) (Nakabachi *et al*., 2022b). This implies that the Secondary_AM/Psyllophila lineage is an ancient symbiont acquired before the divergence of these psyllid genera, although their phylogenetic relationships within the subfamily Psyllinae, and the age of their divergence are uncertain (Burckhardt *et al*., 2021). Secondary_AM and *Psyllophila* showed highly conserved synteny (Fig 3), sharing most genes that exhibit metabolic complementarity with *Carsonella*, including all genes (*trpD, trpCF, trpA,* and *trpB*) required for complementing the incomplete tryptophan biosynthetic pathway of *Carsonella*, and genes for synthesizing B vitamins, biotin (*bioA*, *bioD*, and *bioB*) and riboflavin (*ribA*, *ribD*, *ribB*, and *ribE*), which are completely absent from *Carsonella* (Figs 4 and S2). Conservation of the same set of genes (*trpD, trpCF, trpA,* and *trpB*) for complementing the incomplete tryptophan synthetic pathway of *Carsonella* encoding only *trpE* and *trypG* was also observed in *Enterobacterales* secondary symbionts of *C. eucalypti* (Aphalaridae: Spondyliaspidinae) and *H. cubana* (Psyllidae: Ciriacreminae) (Sloan and Moran, 2012b), which are distantly related to each other and the Secondary_AM/*Psyllophila* lineages (Fig 2). Moreover, genes for synthesizing biotin (*bioA*, *bioD*, and *bioB*) and riboflavin (*ribA*, *ribD*, *ribB*, and *ribE*) were conserved in *Profftella* of *Diaphorina* spp. (Nakabachi, Ueoka, *et al*., 2013; Nakabachi, Piel, *et al*., 2020), which is a versatile symbiont distantly related to the Secondary_AM/*Psyllophila* lineages. These are interesting examples of convergence under selective pressure of limited nutritional availability. Although all the lineages of Secondary_AM, *Psyllophila*, and *Profftella* lacked *ribC* that is required for the final step of riboflavin biosynthesis, previous studies demonstrated that host psyllids horizontally acquired *ribC* from an unknown bacterium before the radiation of the major psyllid lineages (Sloan *et al*., 2014; Nakabachi, 2015; Kwak and Hansen, 2023). Thus, the absence of *ribC* is supposed to be compensated for by this horizontally acquired host gene, showing mutually indispensable tripartite association based on genetic complementarity among the host and symbionts. The involvement of host genes horizontally acquired from bacteria (mostly lineages not leading to extant bacteriome residents) is observed in the bacteriome symbioses of various hemipterans (Nakabachi *et al*., 2005, 2014; Nikoh and Nakabachi, 2009; Nikoh *et al*., 2010; Shigenobu *et al*., 2010; Husnik *et al*., 2013; Luan *et al*., 2015; Mao *et al*., 2018; Ren *et al*., 2020; Smith *et al*., 2021), implying its importance in the evolution of this type of intimate symbiosis. Moreover, newly acquired symbionts repeatedly add or replace the ability to synthesize B vitamins in diverse plant-sap feeding insects (McCutcheon *et al*., 2009; Nakabachi, Ueoka, *et al*., 2013; Husnik and McCutcheon, 2016; Bennett and Mao, 2018; Monnin *et al*., 2020; Nakabachi, Piel, *et al*., 2020; Michalik *et al*., 2021), showing that supplementation of B vitamins is important, if not essential, for these insects (Nakabachi and Ishikawa, 1999). On the other hand, Secondary_AM lacked genes for synthesizing carotenoids (*crtB*, *crtI*, and *crtY*), all of which were retained not only in *Psyllophila* (Dittmer *et al*., 2023) but also in *Profftella* (Nakabachi, Ueoka, *et al*., 2013; Nakabachi, Piel, *et al*., 2020). Carotenoids are organic pigments found in diverse organisms (Fraser and Bramley, 2004; Mussagy *et al*., 2019). In animals, including insects, they are supposed to play important roles, including antioxidation and pigmentation for photoprotection, ornamentation, or camouflage (Fraser and Bramley, 2004; Mussagy *et al*., 2019). Although various microbes and plants produce carotenoids, metazoa cannot generally synthesize carotenoids and must acquire them through diet. Unique exceptions are several arthropod lineages, including aphids, adelgids (both Hemiptera), gall midges (Diptera), and spider mites (Arachnida), which have horizontally acquired carotenoid biosynthesis genes from fungi (Moran and Jarvik, 2010; Altincicek *et al*., 2012; Nováková and Moran, 2012; Cobbs *et al*., 2013). Besides, the genome of “*Candidatus* Portiera aleyrodidarum” (*Gammaproteobacteria*: *Oceanospirillales*), the primary symbiont of whiteflies (Hemiptera: Sternorrhyncha: Aleyrodoidea) encode the same set of carotenoid synthesizing genes (Santos-Garcia *et al*., 2012; Sloan and Moran, 2012a, 2013). As *Portiera* and *Carsonella* are considered sister lineages (Spaulding and von Dohlen, 1998; M. L. Thao and Baumann, 2004; Sloan and Moran, 2012a), their common ancestor may have possessed these genes. However, all *Carsonella* lineages sequenced to date lack them (Nakabachi *et al*., 2006; Sloan and Moran, 2012b; Nakabachi, Ueoka, *et al*., 2013; Nakabachi, Piel, *et al*., 2020; Dittmer *et al*., 2023), suggesting the carotenoid biosynthetic genes of *Psyllophila* and *Profftella* have been complementing the absence of the *Carsonella* orthologs. The lack of these genes in Secondary_AM may reflect the availability of carotenoids in the host plant phloem sap.

Indices of comparisons between genomes of *A. mori* symbionts and their counterparts in *C. melanoneura*, including ANI (Table 2), genome-wide synteny (Figs 3 and S3), similarity of orthologous genes (Tables S1 and S2), and the genome-wide rates of synonymous (*dS*) and nonsynonymous (*dN*) substitutions (Fig 5, Tables S1 and S2), suggested that the Secondary_AM/*Psyllophila* genomes are more labile than those of *Carsonella*. Although its reason is uncertain, the tendency that genomes of younger symbionts are less stable than those of ancient symbionts is widely observed in insect bacteriome symbioses (Bennett *et al*., 2014; Campbell *et al*., 2015; McCutcheon *et al*., 2019; Nakabachi, Piel, *et al*., 2020). This genomic lability may enhance its degradation, leading to the preferential replacement of secondary symbionts, retaining the primary symbionts. Indeed, whereas the Secondary_AM/*Psyllophila* lineage appears to be widely distributed in *Cacopsylla* and some other psyllid genera, several *Cacopsylla* spp. lack the lineage and harbor other secondary symbionts (Nakabachi *et al*., 2022b), suggesting relatively recent replacements. Although some insect lineages appear to have replaced the primary symbiont with newly acquired microbes (Noda, 1977; Fukatsu and Ishikawa, 1996; Sacchi *et al*., 2008; Vogel and Moran, 2013; Nishino *et al*., 2016; Chong and Moran, 2018), it is relatively rare, and no case of *Carsonella* replacement has been identified in Psylloidea.

The genomes of Secondary_AM and *Psyllophila* encoded no genes apparently involved in the defense of the holobiont, highlighting *Profftella*’s uniqueness in having both nutritional and defensive roles. This observation supports the hypothesis that bacteriome-associated secondary symbionts mostly have nutritional roles in complementing the partly degraded genomes of the primary symbionts, as shown in various insect lineages.

## Acknowledgments

We thank Nami Uechi at the Institute for Plant Protection for kindly collecting and providing us with *A. mori* specimens. This work was supported by the Japan Society for the Promotion of Science (https://www.jsps.go.jp) KAKENHI [grant numbers 21687020, 26292174, and 20H02998]. The funders had no role in study design, data collection and analysis, publication decisions, or manuscript preparation.

---

**Fig. S1.**- COG classification of proteins of *Carsonella*_AM and Secondary_AM. The percentage of the total number of genes in each functional category as defined by the COG database for *Carsonella*_AM (green) and Secondary_AM (magenta).

**Fig. S2.**- Amino acid and vitamin biosynthetic genes encoded in the genomes of *Carsonella* and secondary symbionts from *A. mori* (AM), *C. melanoneura* (CM), and *D. citri* (DC). Filled and open boxes indicate the presence and absence of genes, respectively.

**Fig. S3.**- Comparison of the genomic structures of *Carsonella*_AM and *Carsonella* derived from *Cacopsylla melanoneura* (CRmelET). The genomes of *Carsonella*_AM and CRmelET are represented by the x and y axes, respectively. The thick line indicates the shared synteny between the two genomes. The color of the line indicates the percentage similarity between the nucleotide sequences.

## Data Availability

The genomes of Secondary_AM and *Carsonella*_AM have been deposited in DDBJ/EMBL/GenBank databases under the accession numbers AP031380 (*Carsonella*_AM) and AP031381 (Secondary_AM).

