## Supplementary figures and images for "Highly reduced complementary genomes of dual bacterial symbionts in the mulberry psyllid *Anomoneura mori*"

### Fig S1

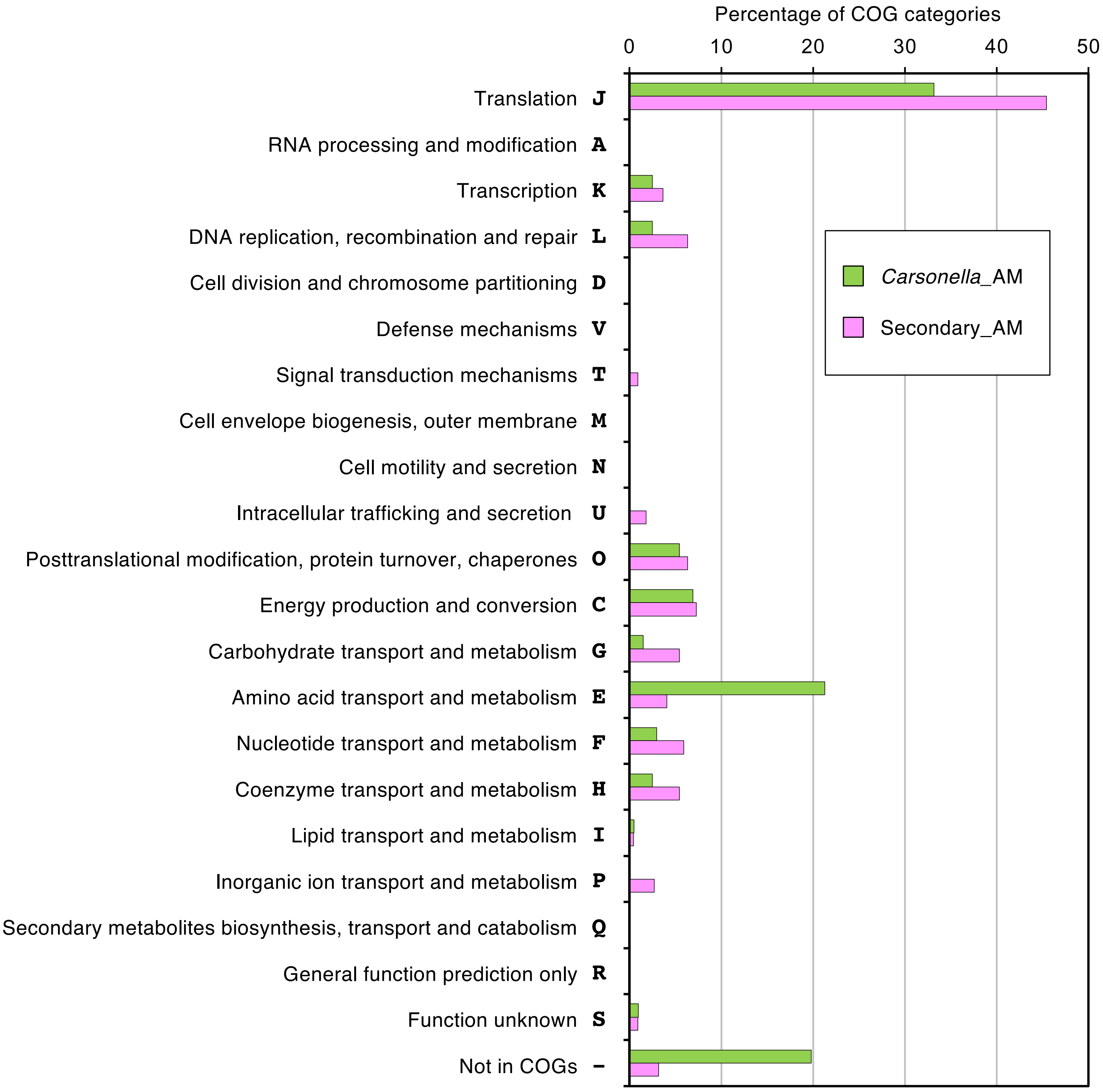

### Fig S2

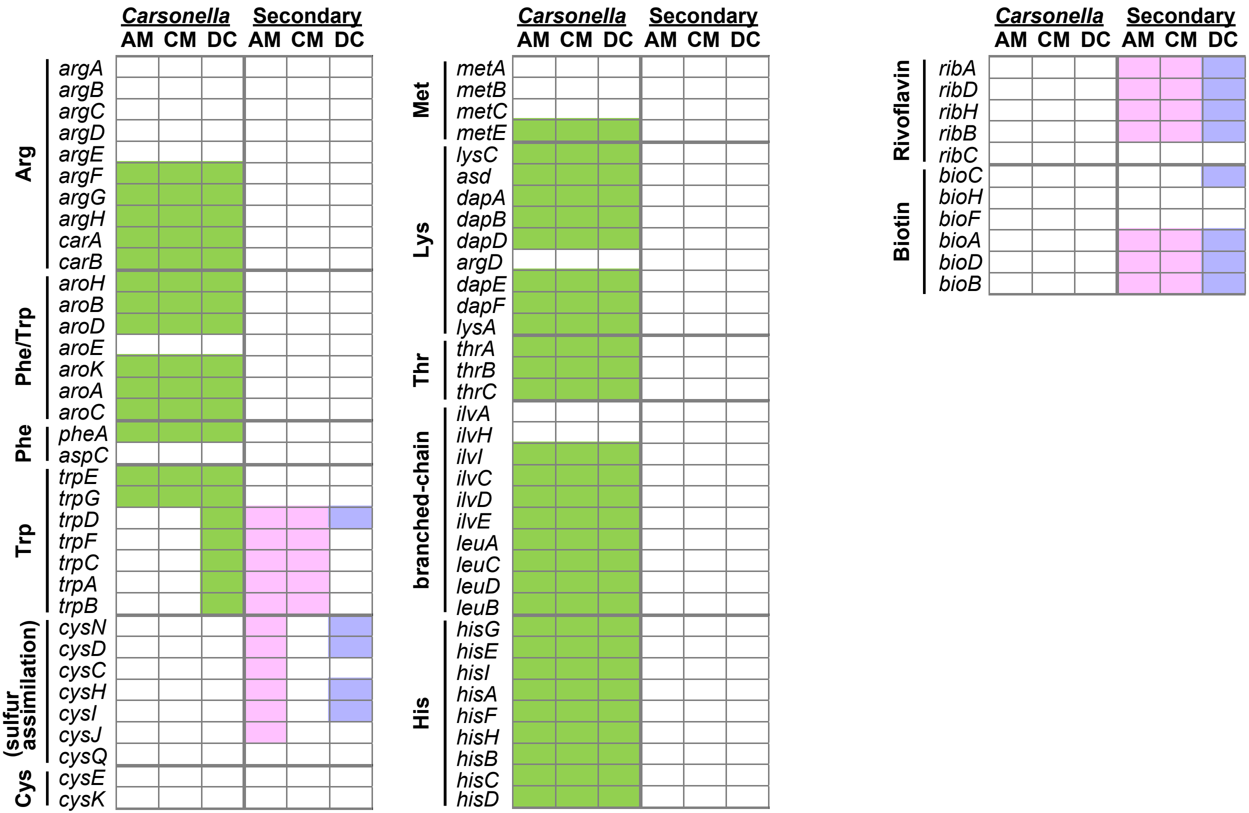

### Fig S3

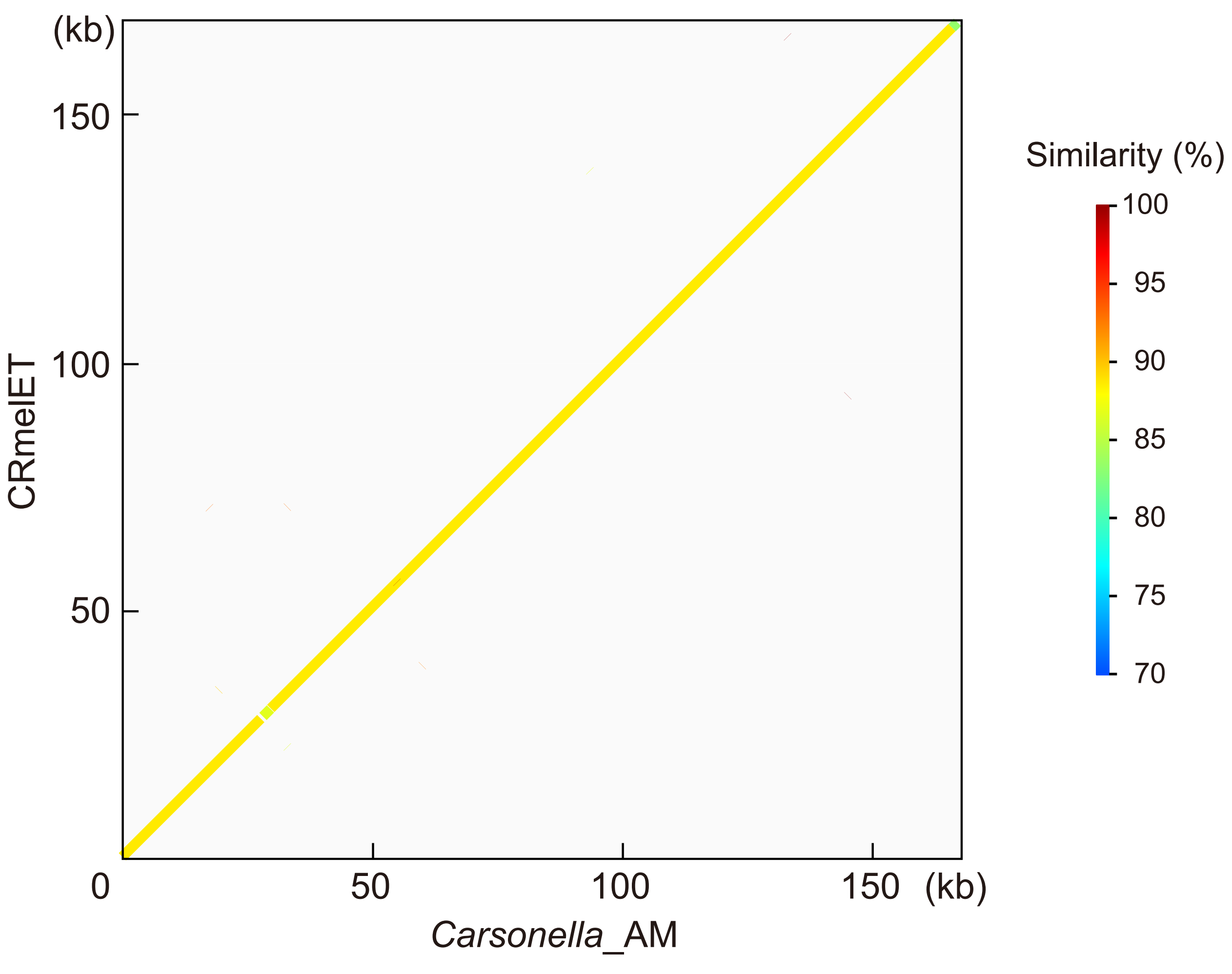
